# From Correlation to Causation: Unraveling the Impact of Closure on Open-Ended Evolution within the Kauffman Model

**DOI:** 10.1101/2024.10.30.621154

**Authors:** M. Faggian

## Abstract

Evolution is a highly intricate process marked by the generation of novelty, which requires the historical and collective organization of individuals. In this paper, we investigate the relationship between biological organization and Open-Ended Evolution (OEE), with a particular focus on the causal connection between the two. To quantitatively investigate the causal relation within a chemical system, we utilize assembly theory to assess the influence of auto-catalytic sets on the complexity dynamics of Kauffman’s model. In the second part of the paper, we strengthen this conjecture by analyzing the effects of the simplest auto-catalytic set on the complexity dynamics in Kauffman’s model, specifically in the absence of parametric correlation. By interpreting auto-catalytic sets as organizational structures in chemical systems, our findings provide the first numerical support for investigating the causal relationship between biological organization and OEE. This work represents a promising area for the initial study of the dynamical relationship between OEE and biological organization, and may advance the understanding of their connection in theoretical biology.

## 1. Introduction

Among the many unresolved questions in mathematics and physics, modeling biological evolution remains a highly complex challenge that has yet to find a unified mathematical framework. At first glance, one might argue that biological evolution does not conform to any of the physical symmetries that we are familiar with. While physical invariances are typically associated with inanimate objects, living systems produce intricate patterns that manifest on a different scale, often within the collective organization of individuals rather than in isolated components.

Classical variables such as time and space, which are essential for describing physical systems, are no longer sufficient to characterize a biological evolution. Instead, the equivalent of time for living organisms is what we refer to as “history.” This shift highlights that space and time, as we understand them, are inadequate for biological systems, further supported by the fact that living beings possess freedom of choice and do not follow predetermined physical trajectories. To properly describe biological evolution, we need to define symmetries that incorporate history and context—two interconnected concepts that influence the evolution of living systems.

One important symmetry in evolutionary biology is biological organization or *Closure* [1, 2, 3], which refers to the specific organization of elements within a living system. When the components of a system meet Closure conditions, they constrain the existence of one another over a time interval. Each element becomes both a constraint on and a product of other constraints, creating a network of interdependencies that drive the system’s organization.

Another key symmetry is the concept of *Open-Ended Evolution* (OEE). Although recent advances in artificial intelligence have provided definitions of Open-Ended Evolution in various contexts [4, 5, 6, 7, 8, 9], these definitions cannot be fully applied to biological OEE. As A. Pocheville noted in [10], *biological sequence may be more complex than its algorithmic counterpart*. In other words, OEE reflects the impossibility of constructing a mathematical description that can encapsulate the evolving system through its history. Models such as the one proposed by Kauffman [11, 12], emphasize the role of self-organization, suggesting a new framework for understanding biological evolution as a boost of historical complexity in a chemical reaction system.

If the ideal mathematical framework for describing evolution remains yet elusive, several efforts have attempted to formalize biological evolution as a transition between two states, based on concepts such as autonomy [13] and undecidability [14, 2].

In the context of chemical systems, several theories have sought to provide a formal framework for understanding evolution in terms of biological organization [13, 15, 16, 17]. If we accept that life’s origins were grounded in chemical reaction systems, then any evolving system that increases its historical complexity must involve a particular form of selection, such as stabilizing, disruptive, or directional selection, which acts as the driving force — “the motor” [18, 2, 19] — behind this increase in complexity. Therefore, natural selection would suffice as the primary force for evolution to emerge in living systems, and evolution must always be linked to the presence of natural selection. In this framework, selection can be seen as the sole physical symmetry responsible for driving evolution in biological systems.

Although modeling the fundamental characteristics of evolution remains a formidable challenge, the Kauffman model of auto-catalysis [20] has provided valuable progress to build the fundaments. Further work by Kauffman [12, 11] has expanded on this interpretation, drawing an analogy between the increase of the historical complexity in auto-catalytic cycles within chemical reactions and the historical complexity augmentation driven by organizational closure in living systems.

By reinforcing the analogy between auto-catalysis and organizational closure, this paper aims to advance the correlation to a causal relationship between the formation of functional organizations in biological systems and the emergence of open-ended evolution, as contextualized within Kauffman’s model [20]. Using assembly theory, we study the evolution of the dynamics of an estimate of the historical complexity of a chemical reaction system within the context of the Kauffman framework. In this analysis, we interpret auto-catalysis as a particular manifestation of organizational Closure in chemical systems. Our investigation delves into the causal influence of auto-catalytic cycles, shedding light on their pivotal role in facilitating the emergence of open-ended evolution in models like Kauffman’s.

## 2. Motivations

As highlighted by various authors [21, 2, 19], understanding OEE mathematically represents an exceptionally challenging problem. The quest for mathematical tools to provide a faithful description of OEE can be framed as the following: *to say everything we can about what we cannot talk about*. In this article, we draw on principles from theoretical biology to present a numerical experiment that attempts to push forward the correlation to a causal relationship between the symmetries emerging within biological evolution. In particular, we provide a numerical study of causal relationship between biological organization as Closure of constraints and Open-Ended Evolution in the setup provided by the Kauffman model.

### 2.1. Closure

As extensively explored in the literature [2, 3], the study of thermodynamic flows within biological systems reveals specific quantities that are conserved across distinct timescales, known as constraints, which represent symmetries inherent in the system [2, 22].

At different timescales, these constraints are subject to degradation and must be either replaced or repaired to maintain the system’s stability. This process can be interpreted differently depending on the scale at which the constraint is being analyzed. For example, while individual enzymes are replaced, their overall population is considered to be repaired [22, 1].

#### 2.1.1. Organization: Closure of constraints

In biological organizations, constraints correspond to physical symmetries that are valid at specific timescales, which may also be interdependent with other constraints. In this paper, we will refer to *Closure* of a set of constraints **C**, when each constraint *C*_*i*_ ∈ **C** directly depends on another constraint *C*_*j*_ ∈ **C** at timescale *τ*_*j*_ (making *C*_*i*_ dependent), and at least one additional constraint *C*_*k*_ ∈ **C** depends on *C*_*i*_ at a different timescale *τ*_*k*_ (making *C*_*i*_ enabling) [2].

Although the mathematical concept of Closure might appear straightforward, its implications from a biological and philosophical perspective are profound [23]. These include the autonomy of living systems and their inherent capacity to self-maintain and self-produce. In this context, Closure enables the system to establish its own goals and norms, while simultaneously promoting the essential conditions for its continued existence through interactions with its environment [1, 2, 22].

An illustrative example of the emergence of organizational Closure is found in Kauffman’s model, where the formation of reflexively auto-catalytic sets of peptides and polypeptides can be associated with a specific realization of Closure [20]. This model offers a simple yet powerful representation of how such closed organizational structures can arise in biological systems.

### 2.2. Principle of Variation

Understanding OEE from a mathematical perspective requires a change in how we perceive the physical world. Specifically, this requires a transformation in the conceptual framework used to model the entities we aim to describe. In physical theories, the primary subjects are referred to as generic objects [21]. These are physical entities of the same kind that follow the same laws. Generic objects are characterized by symmetries that justify the mathematical structure of the phase space and the equations governing their trajectories. These symmetries are time-invariant, meaning they remain unchanged even if the object undergoes some alteration. As such, these symmetries capture the fundamental, unchanging features of the system.

In contrast, biology deals with specific objects [22], which are defined not only by the symmetries that govern generic objects but also by their qualitative characteristics, which are a direct result of their histories. The mathematical description of these specific objects must therefore account for both their symmetries and their adaptation, reflecting the evolution of these symmetries over time.

According to the Principle of Variation [22], which asserts that biological organisms are specific objects, it follows that biological systems undergo unpredictable changes in their symmetries over time. Since these organisms are specific objects, their evolution is influenced not only by their intrinsic symmetries but also by the particular contexts in which they exist and the choices they make—factors that form their unique histories. Because each organism’s context and history are distinct, the mathematical structure representing these objects must evolve unpredictably. Consequently, the way symmetries change in systems of specific objects is not governed by a fixed order, allowing for unpredictable changes at any level of the biological organization.

Systems that adhere to the Principle of Variation in their evolution require reproduction to be sustained in an open-ended manner. The process that drives this ongoing evolution is referred to as Open-Endedness (OEE) or Open-Ended Evolution [22, 24]. This concept highlights the dynamic, unpredictable nature of biological systems, where evolution continues without a predefined endpoint, allowing for continual novelty and adaptation.

#### 2.2.1. Open-Ended Evolution

A recent work by Corominas-Murtra [25] seems to open a venue for putting the basics to a minimal description of OEE. Following briefly their work, if we consider the descriptive state *σ*_*t*_ of a generic system at time *t*, we can define the set 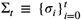 which tracks the *history* of the system at different discrete time steps *t* = 0, 1, … , *t*. They define *K*(Σ_*t*_) the Kolmogorov complexity of the history of the system.

Corominas-Murtra [25] proposed three axioms for the formalization of OEE, but we will focus particularly on the first, which states that the complexity *K*(Σ_*t*_) of an open-ended system does not decrease when divided by time:

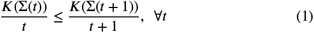

While the increase of complexity alone corresponds to the coming of a novelty in the system, this axiom thresholds the growth of the complexity of an open-ended system at least as the complexity of a random chain.

In continuous limit, it can also be re-written as

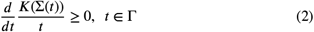

where Γ is a dense time interval.

In other words, if this condition is satisfied then a non-trivial novelty can occur at any time and the system experiences OEE. The non-triviality of the novelty consists in being a functional novelty that changes the organization of the biological system, this it does not only correspond to the discover of a new living entity. The novelty can also be considered as a change of the organization of the constraints of the living system. For example, in a system of chemical reactions the discovery of a new molecule is a novelty in the sense of chemical diversity, but it does not necessarily add a new ysystem functionalit. Thus, the newly discovered molecule might slightly increase the complexity, but not more than a simple random chain. In order for the molecule to increase the complexity in the sense of Corominas-Murtra [25]’s first axiom, it shall induce new functional changes, such as the emergence of a new auto-catalytic cycle in the Kauffman model.

The axioms provided by [25] represent an important contribution to the study of OEE. However, due to the incalculability of the Kolmogorov complexity, it cannot be applied straightforwardly.

If this problem could represent a crucial blocking point in physics, in biology this is not always the case because biological dynamics does not involve generic objects, but specific objects. Exactly because of the nature of specific objects, it follows that the Kolmogorov Complexity might not always be a really appropriate measure of the complexity of biological systems.

In the next section, we describe the mathematical method that we chose to address this question and calculate an estimate of the historical complexity in the Kauffman model.

### 2.3. Measuring Chemical Complexity: Assembly

Recent studies [26, 27] have propelled the mathematical formalization of specific objects, introducing a novel technique to infer the degree of selection within a system using a measure known as Assembly.

The calculation of the Assembly quantity hinges on two key parameters: the assembly index *a*_*i*_, which represents the number of synthesis steps required by the system to produce a particular molecule, and the copy number *n*_*i*_, which denotes the number of copies of a unique molecule present in the system. The term unique here takes on a specific meaning, referring to the concept of a specific object as defined in [22]. In this context, even molecules with identical structures may be considered distinct specific objects due to their different historical contexts.

The Assembly quantity, *A*, is computed [27] as follows:

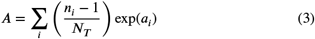

where the sum runs over all unique molecules *i*, and *N*_*T*_ is the total number of molecules in the system.

The authors argue that higher values of this quantity in different realizations signify a higher degree of selection [27].

We propose that this quantity can also be interpreted as a measure of the “effort” involved in constructing the system to reach a certain configuration. As the partition function 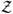 in statistical physics, the Assembly can be viewed as representing the energy required to drive the system to a specific state.

In this article, we employ Assembly not only as a measure of selection within our system but also as an indicator of its complexity. The calculation of Assembly excludes the incidental discovery of new entities (*n*_*i*_ ∼ 1), which contribute minimally to the system’s overall complexity. Thus, it follows that only genuine novelties, defined as new functional organizations within the system, significantly impact its complexity. These new organizations are characterized by the stable production of their constituent entities in non-negligible quantities, which is key to their contribution to the system’s evolving complexity.

## 3. Hypothesis: Closure causes Open-Ended Evolution

In this article, we provide evidence to the hypothesis that not only are functional Closure and Open-Ended Evolution (OEE) strongly correlated, but that Closure serves as the epistemological cause of OEE. As highlighted by [28], while physics currently lacks the tools to provide an explicit causal explanation for physical phenomena, we can interpret the breaking of formal symmetry as being linked to an efficient cause that triggers a change in the system all while preserving its underlying properties.

To illustrate this, consider the electromagnetic force, 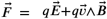. If we focus solely on the electrostatic component 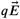 , we can claim that the electric field 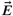 is the cause of the Coulomb force acting on the charge *q*. Similarly, for the magnetostatic component 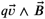, we can say that the magnetic field 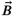 is responsible for the Lorentz force acting on the moving charge.

In biology, the formal symmetry breaking between a simple correlation, denoted by “=“, and a causal epistemic regime is even more significant than in physics [28]. This shift represents a deeper understanding of how complex systems evolve.

To address this issue, we begin with the hypothesis that OEE implies increasing historical complexity, as outlined in Eq. (1) [25]. In a system of chemical reactions, novelty does not simply refer to the discovery of a new chemical species. Rather, it pertains to the novel reorganization of the system’s constraints, which leads to the emergence of a new and unpredictable functional role. Mathematically, this means that while the production of any possible molecule can be predicted at the beginning of time, what cannot be foreseen are the effects that a new organization of constraints will have on the system.

We propose that Assembly serves as an effective measure of the system’s complexity in the context of the Kauffman model. With this formal framework in place, we use Kauffman’s model [20] as a toy model to understand Closure as an auto-catalytic set of chemical reactions. Through numerical analysis, we provide evidence that the emergence of auto-catalysis leads to an increase in complexity within the system, in line with the theoretical insights proposed by [29]. We view Closure and Open-Ended Evolution as two aspects of the same symmetry in biological evolution, akin to the relationship between the electric and magnetic fields in electromagnetism.

Since Closure is represented as a special type of organization of biological constraints, it should only emerge under conditions of strong selection pressure [2, 1]. In contrast, OEE requires the system not only to maintain biological stability across generations, but also to continuously adapt and generate new functional organizations as needed [2, 1, 30].

## 4. Numerical Study: Methods

We examine the evolution of the Kauffman model as described in previous works [20, 29, 31] within the framework of Assembly Theory. In this model, molecules can either undergo ligation, where they bind to another molecule, or cleavage, where they split into two separate molecules. Reactions may be catalyzed with probability *p*_*cat*_ by any chemical species that is not part of the substrate. Catalysis is modeled as a process where the reaction rate is *k*-times faster than that of spontaneous reactions. Under specific conditions, these dynamics give rise to the formation of Auto-Catalytic Sets (RAF) [20].

In this study, we use the term “RAF” to refer to the largest RAF (maxRAF) detected within the system. Similarly, we define the size of the RAF as the total size of the maxRAF, which includes the joint size of all independent RAFs. A more detailed investigation could explore the topological structure of these RAFs, identifying any dependencies of the assembly on the network’s topological properties.

We interpret auto-catalytic networks not merely as a functional novelty but also as a representation of Closure and selection within the system. The molecules that belong to the auto-catalytic network are not only governed by a privileged dynamic but are also selected because their preferential production is essential for the survival and maintenance of the emerging functional organization they form.

To assess the complexity of a chemical system within the Kauffman model, we apply the Assembly Theory, focusing specifically on how the emergence of auto-catalytic networks influences the assembly and overall system complexity.

### 4.1. The Algorithm

We utilize the Gillespie algorithm to run the numerical simulations as in [31].

We consider a chemical reaction system (CRS) defined as in [31] by the tuple 𝒞ℛ𝒮 = (*X, R, C*), where

- *X* is the binary molecules set, in which the most simple molecular subset corresponds to {0, 1};
- *R* is the set of reactions involving the molecules in *X* that are of the type *ligation* when two molecules react to synthesize a new one, or *cleavage* when a molecules splits into two molecules of lower length. Each reaction is exhaustively described by the tuple 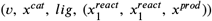 for ligation reactions, and 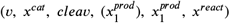 for cleavage reactions, where *v* is^1^the reac^1^tion speed and *x*^*cat*^ ∈ *X* the set of catalysts of the reaction;
- 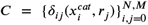 is the set catalytic assignments, where *δ*_*ij*_ is a delta set-selecting function that equals to one if^*ij*^the molecule 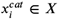 catalyzes the reaction *r*_j_ and to zero otherwise, *N* is the total number of molecules in the system, *M* the total number of reactions, and 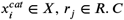 is then a subset of the product set *X* × *R*.

It is important to note that the delta function *δ*_*ij*_ not only associates the catalyst to the reactions, but it also contains information about the topology of the model.

The systems always disposes of an inexhaustible subset of molecules ℱ named *substrate*, which corresponds to the ensemble of all the possible binary molecules with length up to *F* = max_ℱ_ *l*(*x*|*x* ∈ ℱ), where *l* : *x*→ ℝ^+^ is the measure of the length of a molecule *x*. An incoming stream of molecules allows each chemical species belonging to the substrate to never decrease under the initial quantity *Q*_ℱ_ . Only the molecules *x*_*i*_ ∈ *X* with length up to ℕ ∋ *L* = max *l*(*x*|*x* ∈ *X*), *F* < *L* , are allowed to exist in the system. The speed of catalytic reactions is set to be *k*-times the value of the spontaneous reaction speed *v*_0_: *v*_*cat*_ = *kv*_0_. The choice of *K* and the initial amount of substrate *F*_0_ plays a role on the speed of the simulations, but as we will see later, the conceptual results of our study are independent on them.

For our simulations, we set the maximum molecular length *L* = 8 and the maximum substrate length *F* = 2 which corresponds to ℱ = {0, 1, 00, 01, 10, 11}. With this choice of *F* and *L*, the emergence of RAF is independent on these two parameters [20].

The initial configuration of the system at time *t* = 0 is solely provided by the molecules belonging to the substrate set ℱ, and the reaction dynamics is simulated using Gillespie’s algorithm[32]. The molecules belonging to the substrate ℱ do not catalyze any reaction. Every time a new chemical species is discovered during the evolution of the system, it catalyzes an existing reactions with probability *p*_*cat*_. The value of *p*_*cat*_ plays a crucial role on the time required to observe the emergence of a RAF set. [20, 31]

### 4.2. The Molecules in the System

Our numerical simulations use reflecting boundary conditions, i.e. molecules with maximum length *L* can only react via cleavage reactions and cannot ligate to others. Equivalently, all the ligation reactions that would produce a molecule longer than *L* generate a molecule that is highly unstable. We consider these molecules to not be observable in the system, thus for simplicity in the numerical simulations these reactions are not allowed.

Furthermore, we assume the absence structural symmetries among molecules, i.e. molecules represented by different binary sequences correspond to different chemical types.

### 4.3. RAF Detection

In order to detect the RAF emerging in Kauffman’s model, we adopted the algorithm for the auto-catalytic sets generated by a food source (RAF) described by Hordijk et al. in [33] implemented it in Python 3.8.

#### 4.3.1. Assembly Index

When calculating the assembly of the system, before running the simulations, we calculated the assembly indexes *a*_*i*_ of each chemical species via Monte-Carlo simulations including all the possible relations 4 and 5 for a network of reactions with parameters (*F* , *L*).

In particular, the iterative algorithm to determine *a*_*i*_ for ligation reactions is

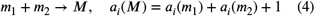

and for cleavage

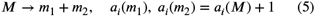

By definition, the assembly index is null for all the molecules belonging to the substrate ℱ. The assembly index is always a positive quantity because the molecules of ℱ do not do cleavage reactions.

In our simulations, we decided to associate the assembly index to a chemical species using what we call *minimal path history of chemical species*: if different assembly indexes are calculated for molecules belonging to the same species, we chose to take the minimum value which corresponds to the shorter assembly path possible in the system [34]. This decision comes from the intent of simplifying the model for numerical purposes, as the distribution of synthesizing paths of a molecular species is very narrow.

There are two more possible ways to calculate the assembly index.

A first alternative approach that we call *history of the chemical species* is to evaluate the the assembly index of each molecule on-the-run, and associate the minimum value to the chemical species whether different assembly paths exist for molecules with the same structure.

A second way that we call *history of the molecules* is to associate to a molecule the index corresponding directly to its assembly path in a numerical simulation. For very large systems and very long simulated times. Both these alternative approaches are equivalent to the aforementioned utilized in our simulations. However, the latter two introduce a stronger historicity in the system that for different types of studies might unveil some more features of evolving biological systems.

### 4.4. Assembly Index in the Simulations

It can be easily shown by induction that a good approximation for the minimum value of the assembly index *a*_*max*_ for uni-dimensional binary molecules in this context is provided by

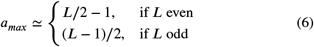

The numerosity Ω(*a*_*i*_) corresponding to the number of chemical species with the same assembly index *a*_*i*_ can be calculated by using Eq.(6) for any value of the molecular length. For example, for even values of the molecular length we get ln Ω(*a*_*i*_) = 2*a*_*i*_ ln 2 + *D*, where *D* ∈ ℝ is a constant that changes depending on *L*.

In our simulations, we found very good agreement for all these quantities, showing that the approximation holds very well. These results are also good evidence for the interpretation of the assembly as historical complexity for a system of chemical reaction of uni-dimensional binary molecules.

## 5. Effects of the Emergence of Auto-Catalysis on the Assembly Dynamics

We consider a network of chemical reactions in which the catalysts 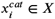 are assigned to the reactions *r*_*j*_ ∈ *R* with constant probability *P* (*r, cat*) :

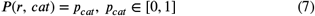

The original formulation of this model was done to ease auto-catalysis-emergence, as can be shown by doing some basic calculations. Indeed, as the probability of associating a catalyst to a reaction *r*_*i*_ is a constant *p*_*cat*_, it follows that the probability that the set of catalysts *X*^*cat*^ of a reaction *r* is not empty is

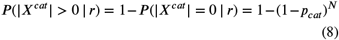

As *N* increases in time, for *t* → ∞ we find

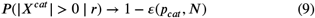

where 1 ≫ *ε* ∈ ℝ ^+^ is a constant that decreases as *p*_*cat*_ increases.

For any *p*_*cat*_ > 0, over extremely long simulation times, it becomes increasingly unlikely for a reaction to remain uncatalyzed indefinitely in an unbounded system. However, this does not always hold for systems of finite size or for numerical simulations conducted over finite time intervals. As demonstrated by [31, 11], the emergence of a RAF (Reflexively Auto-catalytic and Food-generated set) requires the catalytic fraction of reactions to surpass a critical threshold, which depends on the size of the system. In Fig.(1) we report the relation between the numerosity of molecular species versus the assembly index *a*_*i*_, showing that under some hypothesis the assembly index is a good indicator of historical complexity.

**Figure 1:**
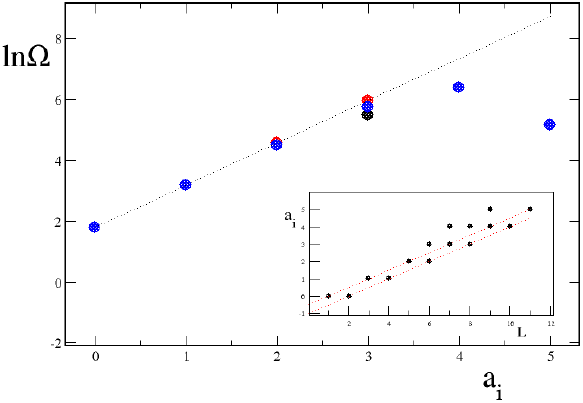
Logarithm of the numerosity of molecular species Ω encountered in the numerical simulations as a function of their assembly index *a*_*i*_. Simulations for *N* = 7 and *F* = 2 (black), for *N* = 8 and *F* = 2 (red) and for *N* = 9 and *F* = 2 (blue). Dashed black line: ln Ω = 2*a*_*i*_ ln 2 + ln 6. Inset: assembly index *a*_*i*_ as a function of the molecular length *L* for different molecular species encountered in the numerical simulations. The two dashed red lines correspond to the two approximations of Eq.(6).

In our study, for *p*_*cat*_ > 0, we primarily focus on values at or below this critical threshold. As demonstrated in the following sections, both the emergence of organizational Closure and the enhancement of assembly are closely linked to *p*_*cat*_. However, this correlation diminishes when operating within a range of very low *p*_*cat*_ values.

### 5.1. Time Evolution of the Assembly

To gain an initial understanding of the temporal behavior of assembly, we performed numerical simulations and measured assembly over time across different realizations of the system for the same value of *p*_*cat*_, as shown in Fig. 2.

**Figure 2:**
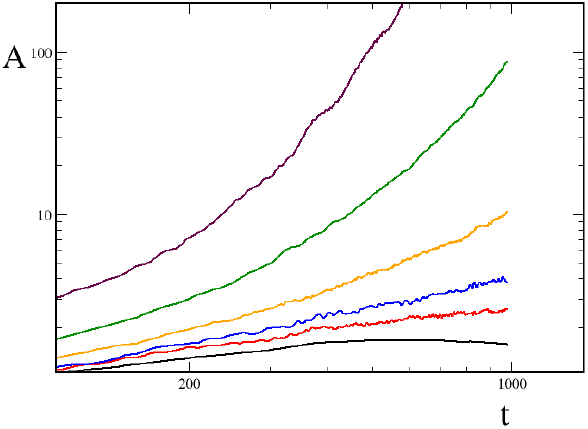
Numerical results of temporal dynamics of the Assembly for different runs of the system with *p*_*cat*_ = 0.01, *Q*_ℱ_ = 10 and *k* = 10^4^ and different RAF sizes at the end of the simulated time: absence of auto-catalysis (black), unitary RAF (red), RAF with final size 3 at the end of the run (blue), final size 10 (yellow), final size 21 (green) and final size 50 (maroon).

The results show that the temporal dynamics of assembly is strongly influenced by the type of RAF present in the system. Specifically, assembly levels are higher in the presence of auto-catalysis, and the rate of change in assembly over time closely correlates with the rate of change in RAF size. In other words, our simulations show that the dynamics of assembly is directly shaped by the dynamics of auto-catalytic cycles. This relationship consistently appears in the Kauffman model when the value of *p*_*cat*_ is sufficiently high to allow the emergence of a RAF.

To further explore the correlation between the emergence of a RAF and enhanced assembly, we apply the initial OEE axiom proposed by [25]. Figure 3 provides a clear example of this axiom: the time derivative of *A*/*t* increases immediately after the RAF emerges, a behavior absent when no RAF is present.

**Figure 3:**
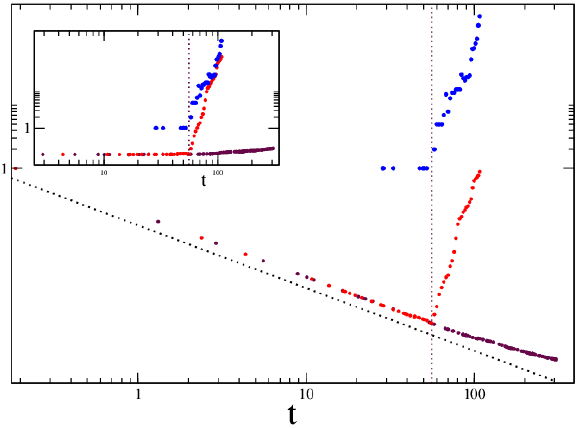
Assembly/time with emergence of auto-catalytic networks (red) and the corresponding curve of the RAF size (blue dots), assembly in absence of RAF (maroon), both the simulations were run for *p*_*cat*_ = 0.002, *Q*_ℱ_ = 10 and *k* = 10^4^. Dashed brown line: 1/*t*. Inset: same data in which the assembly *A* is not normalised by the time *t*.

As noted earlier, the dynamics of assembly is closely tied to *p*_*cat*_. This raises the possibility that the observed behavior simply a result of having a non-null value of *p*_*cat*_. To investigate this further, we first evaluate the correlation between assembly and RAF size, across different *p*_*cat*_ values. In the following sections, we examine the model dynamics in the uncorrelated regime, where *p*_*cat*_ is too low to generate any RAF, specifically when *p*_*cat*_ → 0.

### 5.2. Correlation between Assembly and RAF

To investigate the correlation between assembly and RAF size, we analyzed this relationship across various *p*_*cat*_ values as presented in Fig. 4.

**Figure 4:**
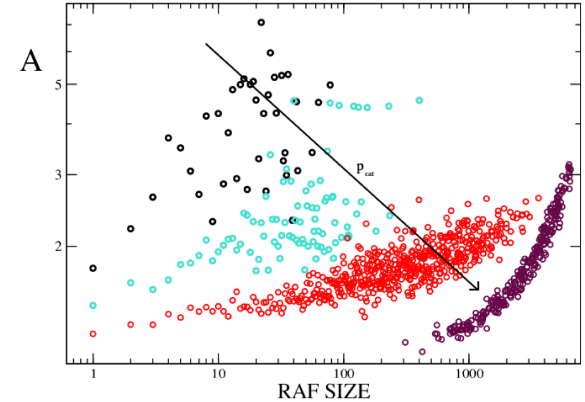
Scatter plot of assembly for *n* = 10 runs as a function of the size of the RAF for *Q*_ℱ_ = 10, *k* = 10^4^ and for different values of *p*_*cat*_: *p*_*cat*_ = 0.003 (black), *p*_*cat*_ = 0.005 (turquoise), *p*_*cat*_ = 0.01 (red) and *p*_*cat*_ = 0.05 (maroon).

If the correlation is now evident, the mere presence of a non-null *p*_*cat*_ prevents us from conclusively stating that auto-catalysis is the direct cause of the increase of the assembly of the system. As detailed in [31], the parameter *p*_*cat*_ is closely correlated not only to the probability threshold for RAF emergence but also to the growth rate of RAFs. Specifically, higher *p*_*cat*_ values increase both the likelihood of RAF formation and the speed of their subsequent growth. To address this complexity, we focus in the following sections on the regime where *p*_*cat*_ ≪ 1, a range in which the assembly dynamics is decoupled from *p*_*cat*_. The central question we aim to answer in this part of the study is : can the smallest RAF induce open-ended dynamics in the Kauffman model?

## 6. Introducing an auto-catalytic molecule: Effects on the Assembly Dynamics

To gain a deeper understanding of the causal relationship between the emergence of RAFs and the increase in complexity within the Kauffman model, we examine the effect of a minimal-size RAF on assembly dynamics. The introduction of such a molecule represents a deviation from the original version of the Kauffamn model, as the model was initially conceived in such a way that no molecule need catalyze its own formation.

Specifically, we investigate the model’s behavior in the limit *p*_*cat*_ → 0, analyzing the impact of introducing a unitary-sized RAF (URAF), also known in literature as *auto-catalytic molecule*, compared to a simple catalytic reaction (CR). In the simulation, both the URAF and the CR are activated when their respective catalyst is introduced into the system at time *t*_0_, with the catalyst acting as a “seed” for complexity growth. If organizational Closure is indeed responsible for the assembly boost, we expect the URAF to influence the dynamics in a fundamentally different way than a simple CR.

To quantify this, we compare the growth of complexity in two systems—one with a URAF and the other with a CR—using the first axiom from [25].

### 6.1. Introduction of an auto-catalytic molecule

When in the presence of a URAF, the time behavior of the assembly is pretty simple to understand.

*A*_0_ is the assembly contribute of the catalyst *C*, called the *seed* of the system. One could express the total assembly of the system as the sum of several contributions:

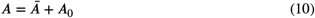

where

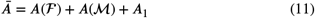

where *A*(ℱ) is the assembly of the substrate (i.e., the molecules with assembly index equal to zero), *A*(ℳ) the contribution of the molecules ℳ that have *C* as common ancestor in the chain of reactions, and *A*_1_ the contribution of all the other molecules.

The contribution of the molecules in *A*(ℱ) is negligible over time, as their assembly index is null. The amount of substrate ℱ can be considered almost constant over time: *A*(ℱ)/*t* → 0. The contribution *A*_1_ is negligible, as the molecules that contribute to this are synthesized by spontaneous processes. Thus, we can make the following approximation:

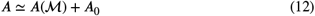

Using a biological metaphor, we can think of the assembly as being shaped by two main contributors: the seed-contribute *A*_0_, and the components of the system that grow around the seed, *A*(ℳ). While the seed-contribute serves as the system’s fuel over an extended period, its influence gradually diminishes in the long run, allowing the dynamics of *A*(ℳ) to dominate. When the derivative of the dominant contributor *A*(ℳ)/*t* is non-negative, we interpret this as a sign of open-ended evolution (OEE). In our case, the introduction of a URAF into the system drives the force behind the Open-Ended dynamics.

#### 6.1.1. Numerical results

We developed an algorithm that introduces an auto-catalytic network of a given size *L*_*RAF*_ during a numerical realization of the Kauffman model’s dynamics for *p*_*cat*_ = 0.0, with the objective of studying the impact of such imposition on the assembly dynamics.

We anticipate that introducing a URAF will enhance the system’s assembly, driving it to achieve a higher level of complexity in its dynamics compared to a simple catalytic set (CR). We examine a chemical reaction of the type:

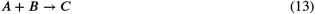

in which the couple *A, B* ∈ ℱ and *C* ∉ ℱ. For the URAF case, we consider the reaction catalyzed by *C*, whereas for a CR it is catalyzed by another molecule *D* ∉ ℱ, to avoid constant catalysis due to the permanent presence of the substrate.

To offer some analytical insights, we examine the time evolution of two types of assembly in the case of a URAF: the total assembly of the system *A*(*t*) and the complementary assembly *Ā*(*t*), which represents the assembly of the system excluding the catalyst.

For both the URAF and the CR scenarios, we consider the same reaction type: a ligation-type reaction involving two different molecules from the substrateℱ. In Fig. 5, we show the temporal behavior of the total assembly for both the URAF and the simple CR cases, along with the time evolution of *Ā*(*t*) after introducing the URAF.

**Figure 5:**
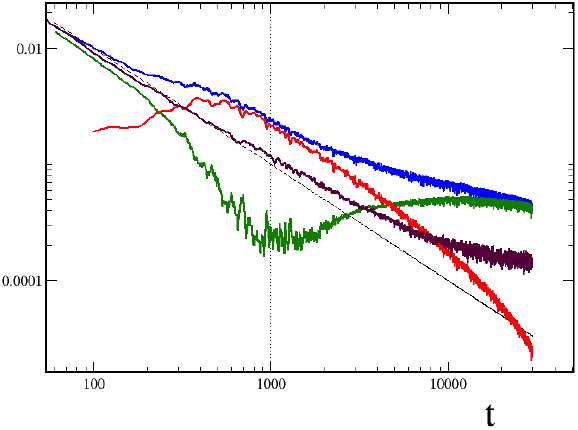
Time evolution of the assembly with URAF compared to the assembly with CR, for *p*_*cat*_ = 0.0, *Q*_ℱ_ = 10, *k* = 10^4^ and *t*_0_ = 0.0. The curves represent: *A*(*URAF* )/*t* (Blue), *A*(*CR*)/*t* (maroon), *A*_0_/*t* (red), (*A*−*A*_0_)/*t* (green). Curves averaged over *n* = 50 realizations.

The numerical simulations reveal that the assembly in the system with a URAF is consistently higher than in the CR case. Additionally, the URAF induces a distinct dynamic in the model compared to the scenario without auto-catalysis. Specifically, the URAF triggers a wave of complexity that increases the likelihood of the system discovering new complex molecules. Furthermore, the URAF acts as a fuel for other chemical reactions, accelerating the system’s discovery of more chemical species. This process continues until the number of molecules in *A*−*A*_0_ becomes negligible compared to the quantity of *C*.

This result is significant. When we subtract the contribution of the exponentially growing URAF to the assembly, the remaining complexity dynamics follow a non-decreasing trajectory, demonstrating open-ended evolution over a substantial time interval. This is clearly illustrated in Fig. 5, where the green curve represents this non-decreasing dynamics after the URAF has saturated the system. The curve shows a continuous growth, driven by a cascade of increasingly complex molecules initiated by the URAF’s initial exponential growth. Eventually, the curve reaches a quasi-plateau due to finite size effects.

#### 6.1.2. Integration of kinematic equations

In order to fully understand the dynamics of the Kauffman model from a perspective of assembly theory, we also studied the kinematic equations for a URAF inserted at *t*_0_ = 0.0 The types of molecules present in the system divide in four principal categories: the molecules belonging to the food *N*_*F*_ , the molecule generated by the URAF and denoted with *N*_*C*_ , the molecules generated using reactions that involve the catalyst of the nucleus *C* that we call *N*_*MC*_ , and finally all the other molecules that are generated by reactions that involve molecules of the food and that do not belong to the ensemble of *MC*, that we call *N*_*MF*_ .

In equations, *N*_*T*_ = *N*_*F*_ + *N*_*C*_ + *N*_*MC*_ + *N*_*MF*_ . Let’s consider the kinematic equations of the chemical reactions involving all these molecules. The auto-catalytic reaction that produces the molecule *C* can be written as 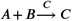, where *A* ≠ *B*, and both belong to the food set (*A, B* belong to a subset of food that corresponds to a fraction 1/6 of the total initial molecules). Defining the quantity *Q* as the initial quantity for each molecule type in the food set, *v* the non-catalytic reaction speed, *k* the ration between the catalytic and the non-catalytic speed, and denoting with *n* the numerical density of a molecule in the system:

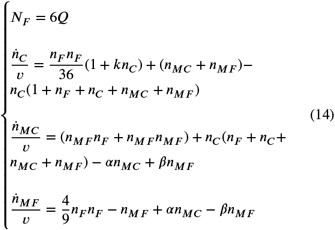

where 0 < *α* < 1 corresponds to the fraction of reactions that from the set *MC* create a molecules that belongs to *MF* , and 0 < *β* < 1 to the fraction of reactions from *MF* to *MC* These two coefficients represent the common molecules between *MF* and *MC*.

Following these equations, *n*_*MF*_ becomes relevant for long time intervals and in particular we find *n*_*MF*_ ∼ *αn*_*MC*_ ).

A good estimate of the parameters *α* and *β* allows to better predict the time behavior of the molecules of type MC. Fig.(6) are reported the results of the numerical integration of such equations that show a good compatibility with the behaviors shown in Fig.(5). In particular, the kinetic equations allow one to better appreciate the presence of a time-asymptotic plateau for (*A* − *A*_0_)/*t*, showing that the presence of a URAF in the system boosts its creativity and pushing its evolution to a non-negative time derivative. The differences between the time behavior of (*A* − *A*_0_)/*t* in the kinematical simulations and the Gillespie simulations are due to a imperfect estimate of the parameters *α* and *β*.

**Figure 6:**
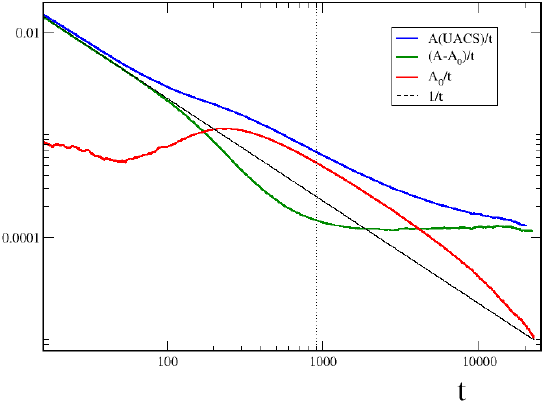
Numerical simulations of the differential equations of kinematics for the total Assembly A of the system and the Assembly of the URAF *A*_0_, for *p*_*cat*_ = 0.0, *Q*_ℱ_ = 10, *k* = 10^4^ and *t*_0_ = 0.0. The different curves use the same color scheme as in Fig.(5) and represent: *A*/*t* (red),*A*_0_/*t* (blue), (*A* − *A*_0_)/*t* (green). Curves averaged over *n* = 50 realizations.

Finally, we want to point out that one might consider controversial the finding of OEE in this set of kinematic equations, because, as we said in the beginning of this article, an open-ended evolving system is not supposed to be described by a finite number of equations[22]. We will return to this point in the conclusions of the article.

#### 6.1.3. The characteristic time τ

By defining *τ* as the value of *t* at which the time-derivative of *Ā*/*t* is not anymore negative and the second time-derivative changes sign (see Fig.(5)), we can define two main regimes for the total assembly *A*: for *t* < *τ* we find *A* ∼ *A*_0_, whereas for *t* ≫ *τ* we have *A* ∼ *Ā*. We explain this behavior with the kinetic equations. For *t* < *τ*, the dynamics of the number of seeds obeys the approximate differential equation 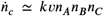 where we identify with *n* the density of molecules in the system.

By solving the differential equation, we find that the number of seeds in the system follows an exponential growth of the type *N*_*c*_ ∼ exp(*kvQ*_ℱ_ *t*) ≡ exp(*t*/*τ*), which slows down when the number of seeds saturates the system for finite-size effects around *t* ≃ *τ*, and is followed by a cascade of reactions involving more complex molecules. In the second phase, for *t* ≫ *τ* the growth rate of the seeds is almost constant, bringing *N*_*c*_ to grow linearly, *n*_*c*_ ∼ *kvQ*_ℱ_ *t*. In this regime, due to the competition between the presence of *C* and that of its descendants, the main contribution of the total assembly *A* is provided by moleculesℳ. This is equivalent to stating that the dynamics presents a complexity wave of molecular length boosted by the initial URAF.

Considering *A*_0_ ∼ *en*_*c*_ ≡ *eN*_*c*_/*N*_*T*_ and approximating the solution of the differential equation of *n*_*c*_ in Eq(14), we obtain the following:

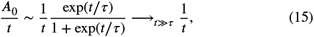

which explains the convergence *A* → *A*_0_ for *t* < *τ*.

Concerning the link between *τ* and the system size *L*, as reported in Fig.(7) the numerical simulations confirm that the value of *τ* converges to *τ* ≃ 1/*kvQ*_ℱ_ for *N* > 5. This confirms the independence of our findings on the parameters *k* and *Q*, showing an adaptation of the dynamics on different timescales.

**Figure 7:**
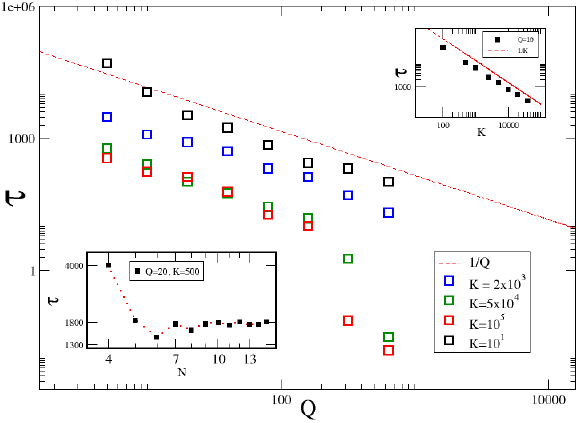
Measurements of *τ* in the realization of the Kauffman model for different values of the initial amount of *Q*_ℱ_ and *k*. The results are in agreement with *τ ∼* 1/*Q*. Inset bottom left: dependency of *τ* on the max length parameter *N*. The value of *τ* converges for increasing values of *N*. Inset top right: dependency of *τ* on *k*. The results show a good agreement with *τ ∼* 1/*k*

#### 6.1.4. Impact of τ on the removal of a URAF

We also studied the case of the removal of a URAF at *t* > *t*_0_, which showed that the time-derivative of *A*(ℳ)/*t* does not become positive at any moment *t* if the seed is removed at *t*_0_ < *τ*. If instead the seed is removed at *t*_0_ ≥ *τ*, the dynamics remains as it is in Fig.(5). Indeed, for *t* > *τ* the assembly dynamics does not depend anymore on the presence of the catalytic seed, and the boost of complexity depends only on the inertial complexity wave of chemical reactions boosted by the initial URAF. In other words, for *t* > *τ* the removal of the URAF does not seem to have an impact on the assembly dynamics, because the same dynamics seems to have started to follow an Open-Ended Evolution path.

### 6.2. Long time behavior

For what concerns the behavior for *t* ≫ *τ*, we find that for *t* > *τ* the value of *Ā* reaches a long temporal plateau after easing for a ver long time interval. We call *Ā* the value *Ā* on this plateau. Fig(8) shows the *N*-scaling of *Ā* _∞_ for *t* ≫ *τ*.

**Figure 8:**
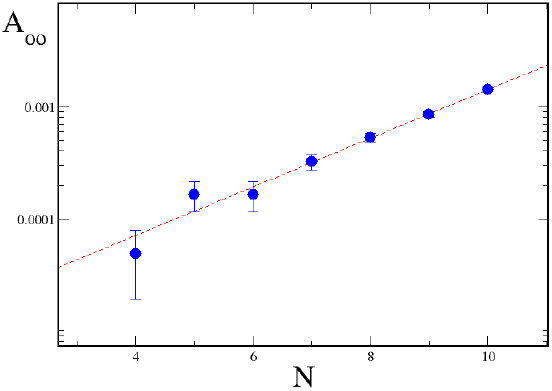
Values of *Ā*_∞_ for *F*_0_ = 10 and *K* = 10^4^ (blue dots). Red dashed line: *Ā*_∞_ *∼* exp(*N*/2).

This dependence on *N* is a direct consequence of the definition of *A*(ℳ). In particular, one can show that *A*(ℳ) is dominated by the terms with higher assembly index:

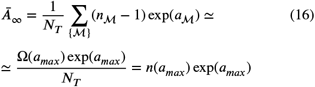

where Ω(*a*_*max*_) is the total number of molecules with assembly index *a*_*max*_ and *n*(*a*_*max*_) their density.

By using the approximation of Eq.(6), we get

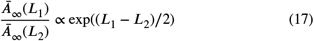

which explains well the results reported in Fig(8), showing an evidence for unbounded OEE in absence of finite-size effects for *L* → ∞

### 6.3. Imposition of a URAF at different *t*_0_

Lastly, we decided to study the effects of the introduction of a URAF at times different than *t*_0_ = 0 to check whether the dynamics studied in the previous sections does or not depend on the configuration of the system.

As shown in Fig. 9, we observed that the system’s reaction after introducing a URAF is independent of the value of *t*_0_, indicating that the complexity boost in the Kauffman model is not influenced by the time at which it begins. This also demonstrates the absence of nucleation in our specific setup.

**Figure 9:**
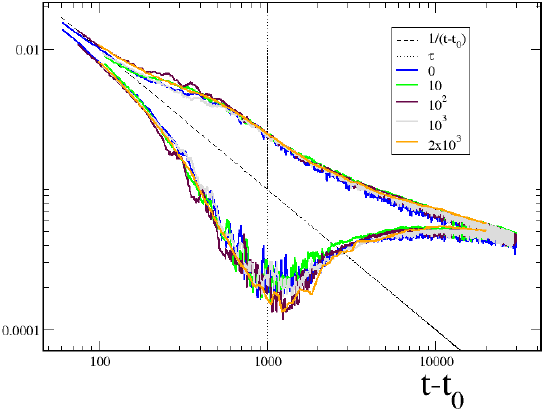
Curves of total assembly *A*/*t*) (top) and *Ā*/*t* (bottom) for different values of the time of introduction of the URAF *t*_0_ reported in the legend box. Curves averaged over *n* = 50 realizations.

## 7. Discussion and Further Developments

We investigated the impact of the emergence of an auto-catalytic network on the dynamics of the assembly in the Kauffman model. Auto-catalysis was interpreted as the simplest representation of organizational Closure in a system, while assembly served as a measure of its complexity. We examined the relationship between the size of auto-catalytic cycles and assembly, showing how cycles of varying sizes influence the assembly. Our findings reveal that the onset of auto-catalysis is a pivotal event in assembly dynamics, as even the smallest cycle induces significant changes in the behavior of the chemical system. Specifically, numerical simulations indicate that the mere presence of a minimal auto-catalytic cycle is sufficient to trigger the emergence of open-ended assembly dynamics, as defined by [25], regardless of the model parameters. This pattern was consistently observed in the Kauffman model under various conditions and parameter choices. Whenever open-ended behavior emerged, the presence of auto-catalysis was the only invariant.

These findings, derived from analyzing the dynamics of the Kauffman model, could be further explored by examining different variations of the model. One potential variation involves allowing *p*_*cat*_ to adapt dynamically, maintaining either *P* (*r, cat*) or the system’s catalytic fraction at a constant level. Another possible variation introduces a pro/anti ancestor bias on *p*_*cat*_, where the likelihood of a molecule acting as a catalyst depends on whether at least one of the reactants in the reaction is an ancestor.

While this study focused on the simplest version of the Kauffman model, the same results are expected to hold across these alternative model variations.

From the perspective of nucleation theory, another promising avenue of research involves examining the dynamics of auto-catalytic cycles in detail for non-zero values of *p*_*cat*_. We hypothesize that the nucleation of the auto-catalytic core will significantly accelerate assembly dynamics, surpassing the simple plateau observed when *p*_*cat*_ = 0. Specifically, we anticipate that during nucleation, the system will exhibit a non-zero derivative of *A*/*t* for non-zero *p*_*cat*_. Additionally, it is crucial to investigate the impact of the topology of auto-catalytic cycles and their sub-cycles on the increase in complexity. For instance, we expect that more interconnected cycles will exert a greater catalytic effect, while cycles with simpler topologies may also demonstrate a higher efficiency. Finally, another research direction involves refining the measure of complexity for systems like Kauffman’s. A more specific definition of assembly tailored to binary molecules could better capture the nuances of open-ended evolutionary dynamics in such models. For shorter binary molecules, the assembly index effectively reflects system historicity, but this approximation may break down for longer non-binary molecules, such as DNA. For instance, redefining the assembly index to account for the compressed size of each molecule could unveil additional features, particularly in the context of multidimensional non-binary molecular systems.

## 8. Conclusions

In this study, we explored the emergence of open-ended evolution (OEE) within the simple framework of the Kauffman model. Using assembly theory, we investigated the influence of auto-catalytic cycles on the complexity dynamics. Our findings offer the first numerical support for a hypothetical causal relationship between organizational closure—exemplified by autocatalysis—and open-ended evolution (OEE), when system assembly is interpreted as a measure of historical complexity.

To further develop this hypothesis, we examined the dynamics of the system under the condition *p*_*cat*_ = 0, introducing a minimal auto-catalytic cycle. Our results show that even the smallest auto-catalytic cycle is sufficient to enhance the system’s complexity, enabling OEE to persist over extended time scales. Furthermore, we observed that nucleation does not occur in the system for *p*_*cat*_=0, although it is expected to arise for*p*_*cat*_ > 0.

Our contributions can be summarized as follows:

- Theoretical Insight: we offer numerical support for a hypothetical causal link between organizational Closure and Open-Ended Evolution in chemical systems.
- Qualitative Analysis: we applied a novel approach for examining the impact of the emergence of auto-catalysis in chemical reaction systems.
- Application of Assembly Theory: we showed a new use of assembly theory as a tool to study the dynamics of complexity in theoretical biology.

While this work provides a promising foundation for understanding biological evolution, further research is essential to bridge the gap between our current knowledge of physical systems and biological entities. Life’s elusive nature has long challenged scientific inquiry, but the principles of physics—particularly the symmetries imprinted by life’s evolutionary history—may ultimately hold the key to unlocking its mysteries.

## 9. Statement

The author claims to have developed the content of this paper independently without taking inspiration from others.

## 10. Acknowledgements

The author thanks A. Pocheville, M. Barbier and G. Folena for prolific scientific discussions and Yoshi D.F. for insightful conversations.

This project/ publication was made possible through the support of Grant 62220 from the John Templeton Foundation. The opinions expressed in this publication are those of the author(s) and do not necessarily reflect the views of the John Templeton Foundation.

## References

[1] Matteo Mossio, Maël Montévil, and Giuseppe Longo. Theoretical principles for biology: Organization. Progress in Biophysics and Molecular Biology, 122(1):24–35, October 2016.

[2] Alvaro Moreno and Matteo Mossio. Biological autonomy: a philosophical and theoretical enquiry, volume 12. Springer, 2015.

[3] Maël Montévil and Matteo Mossio. Biological organisation as closure of constraints. Journal of Theoretical Biology, 372:179– 191, may 2015. https://montevil.org/publications/articles/2015-MM-Organisation-Closure-Constraints/.

[4] Alyssa Adams, Hector Zenil, Paul C. W. Davies, and Sara Imari Walker. Formal Definitions of Unbounded Evolution and Innovation Reveal Universal Mechanisms for Open-Ended Evolution in Dynamical Systems. Scientific Reports, 7, April 2017.

[5] Pedro A. Castillo and Juan Luis Jiménez Laredo, editors. Applications of Evolutionary Computation: 24th International Conference, EvoApplications 2021, Held as Part of EvoStar 2021, Virtual Event, April 7–9, 2021, Proceedings, volume 12694 of Lecture Notes in Computer Science. Springer International Publishing, Cham, 2021.

[6] Department of EECS (Computer Science Division) University of Central Florida, Orlando, FL 32816, L. Soros, and Kenneth Stanley. Identifying Necessary Conditions for Open-Ended Evolution through the Artificial Life World of Chromaria. In Artificial Life 14: Proceedings of the Fourteenth International Conference on the Synthesis and Simulation of Living Systems, pages 793–800. The MIT Press, July 2014.

[7] Nicolas Loeuille, Michel Loreau, and Simon A. Levin. Evolutionary Emergence of Size-Structured Food Webs. Proceedings of the National Academy of Sciences of the United States of America, 102(16):5761–5766, 2005.

[8] Hiroki Sayama. Seeking open-ended evolution in Swarm Chemistry. In 2011 IEEE Symposium on Artificial Life (ALIFE), pages 186–193, Paris, France, April 2011. IEEE.

[9] Susan Stepney. Modelling and measuring open-endedness. Artificial Life, 25(1):9, 2021.

[10] Arnaud Pocheville. Biological information as choice and construction. Philosophy of Science, 85(5), July 2018.

[11] Stuart A. Kauffman. The Origins of Order: Self-organization and Selection in Evolution. Oxford University Press, 1993. Google-Books-ID: lZcSpRJz0dgC.

[12] Stuart A. Kauffman. Antichaos and Adaptation. Scientific American, 265(2):78–84, August 1991.

[13] Kepa Ruiz-Mirazo, Juli Peretó, and Alvaro Moreno. A Universal Definition of Life: Autonomy and Open-Ended Evolution.

[14] Troy Day. Computability, Gödel’s incompleteness theorem, and an inherent limit on the predictability of evolution. Journal of The Royal Society Interface, 9(69):624–639, April 2012.

[15] Kepa Ruiz-Mirazo and Alvaro Moreno. Basic autonomy as a fundamental step in the synthesis of life. Artif. Life, 10(3):235–259, 2004.

[16] Alvaro Moreno and Arantza Etxeberria. Agency in natural and artificial systems. Artif. Life, 11(1-2):161–175, 2005.

[17] Kepa Ruiz-Mirazo and Alvaro Moreno. Autonomy in evolution: from minimal to complex life. Synthese, 185(1):21–52, March 2012.

[18] Börje Ekstig. Complexity, Natural Selection and the Evolution of Life and Humans. Foundations of Science, 20(2):175–187, June 2015.

[19] Francis Bailly and Giuseppe Longo. BIOLOGICAL ORGANIZATION AND ANTI-ENTROPY. Journal of Biological Systems, 17(01):63–96, March 2009.

[20] Stuart A. Kauffman. Autocatalytic Sets of Proteins. J. theor. Biol., 119:1–24, 1986.

[21] Longo G. (2011) Bailly, F. Mathematics and the natural sciences; the physical singularity of life. volume 16, pages 309–336. 2011.

[22] Maël Montévil, Matteo Mossio, Arnaud Pocheville, and Giuseppe Longo. Theoretical principles for biology: Variation. Progress in Biophysics and Molecular Biology, 122(1):36–50, October 2016.

[23] H. R. Maturana and F. J. Varela. Autopoiesis and Cognition: The Realization of the Living. Springer Science & Business Media, December 2012. Google-Books-ID: iOjVBQAAQBAJ.BioSystems

[24] Arnaud Pocheville. A Darwinian dream: on time, levels, and processes in evolution. In Tobias Uller and Kevin N. Laland, editors, Evolutionary Causation. Biological and philosophical reflections, Vienna Series in Theoretical Biology. MIT Press, Boston, 2019.

[25] Bernat Corominas-Murtra, Luís F. Seoane, and Ricard Solé. Zipf’s Law, unbounded complexity and open-ended evolution. Journal of The Royal Society Interface, 15(149):20180395, December 2018.

[26] Stuart M. Marshall, Douglas G. Moore, Alastair R. G. Murray, Sara I. Walker, and Leroy Cronin. Formalising the Pathways to Life Using Assembly Spaces. Entropy, 24(7):884, June 2022.

[27] Abhishek Sharma, Dániel Czégel, Michael Lachmann, Christopher P. Kempes, Sara I. Walker, and Leroy Cronin. Assembly theory explains and quantifies selection and evolution. Nature, 622(7982):321–328, October 2023.

[28] Longo G. (2006) Bailly, F. Mathématiques et sciences de la nature. la singularité du vivant. 2006.

[29] Wim Hordijk and Mike Steel. Autocatalytic Networks at the Basis of Life’s Origin and Organization. Life, 8(4):62, December 2018.

[30] Arend Hintze. Open-Endedness for the Sake of Open-Endedness. Artificial Life, 25(2):198–206, May 2019.

[31] Wim Hordijk and Mike Steel. Chasing the tail: The emergence of autocatalytic networks. Biosystems, 152:1–10, February 2017.

[32] Daniel T Gillespie. A general method for numerically simulating the stochastic time evolution of coupled chemical reactions. Journal of Computational Physics, 22(4):403–434, December 1976.

[33] Wim Hordijk, Joshua I Smith, and Mike Steel. Algorithms for detecting and analysing autocatalytic sets. Algorithms for Molecular Biology, 10(1):December 2015.

[34] Keith Y Patarroyo, Sara I Walker, Abhishek Sharma, and Leroy Cronin. AssemblyCA: A Benchmark of Open-Endedness for Discrete Cellular Automata.

